# Transcriptome-wide analysis of auxin-induced carotenoid accumulation in *Chlorella* microalgae

**DOI:** 10.1101/334102

**Authors:** Faisal Alsenani, Taylor J. Wass, Ruijuan Ma, Eladl Eltanahy, Michael E. Netzel, Peer M. Schenk

**Author notes:** FA and TJW contributed equally to this study. Email addresses.

## Abstract

Microalgae are a commercially viable route for the production of carotenoids, including β-carotene and astaxanthin. In the current study, the commercially relevant microalga, *Chlorella* sp. BR2 was treated with four plant hormones: indole-3-acetic acid, salicylic acid, abscisic acid and methyl jasmonate, over a range of dosages and screened for enhanced carotenoid production. Indole-3-acetic acid was the only hormone with an inductive effect on carotenoid accumulation. As such, the transcriptome under the condition with the highest carotenoid increase was profiled using RNA-Seq and expressed sequences reconstructed with *de novo* assembly. This allowed for the profiling of transcriptome-wide changes following auxin treatment, revealing the active pathway components of auxin-induced carotenogenesis. Data analysis specified the differentially expressed genes involved in auxin biosynthesis and signal transduction, which suggest a close relationship to equivalent pathways in higher plants. However unlike in plants, the ancient ABP1/SCF^SKP2A^/IBR5-mediated pathways for auxin response likely acted as the primary signaling route in *Chlorella*. As carotenoids are precursors for abscisic acid, the findings suggest a causative link between auxin signaling and abiotic stress tolerance.

**Highlight:** Transcriptomics of plant hormone-treated *Chlorella* revealed the active pathway components of auxin-induced carotenogenesis and included the ancient ABP1/SCF^SKP2A^/IBR5-mediated pathways. The manuscript presents the first documented transcriptomic data of auxin-treated microalgae.

## Introduction

Among the 600 carotenoid compounds in nature, only 40 form part of the human diet (Vershinin 1999). Carotenoids such as β-carotene and α-carotene are mainly found in carrots, pumpkin and squash, while lutein and zeaxanthin are found in dark-green leafy vegetables, and lycopene is delivered mostly by tomatoes (Jha et al. 2010).

Microalgae provide large amounts of β-carotene and have been extensively studied to be an alternative source of various other carotenoids, due to their metabolic diversity and fast growing characteristics (Guedes et al. 2011). Some additional carotenoids have already been commercialized, such as astaxanthin from *Haematococcus pluvialis*, which was previously harvested unsustainably from salmon stocks (Panel 2005). Carotenoids have a preventive effect on certain major diseases, including eye diseases, such as macular degeneration, in addition to muscular dystrophy, rheumatoid arthritis and cardiovascular ailments (Hudek et al. 2014). Carotenoids may play fundamental roles in cancer prevention due to their antioxidant activity (Ibañez and Cifuentes 2013). Moreover, it has been reported that carotenoids have a protective effect against UV-induced damage to human skin (Stahl and Sies 2012).

Plant hormones have found applications in the cultivation of microalgae. The application of hormones, including abscisic acid (ABA), indole-3-acetic acid (IAA), methyl jasmonate (MJ) and salicylic acid (SA) has been shown to enhance either carotenoid production and/or total productivity across several microalgal species (Tarakhovskaya et al. 2007). Despite the presence of this useful secondary metabolite stimulation process, the underlying changes in gene expression that drive these phenotypes have not been studied at a systems-level in algae. *Chlorella* and most other unicellular photosynthetic microalgae share a common ancestor with multicellular plants, thereby making them powerful model organisms to investigate hormone signaling at a cellular level.

In the current study, *Chlorella* sp. BR2, was treated with plant hormones, IAA, SA, ABA and MJ. Over the range of dosages screened for enhanced carotenoid production, IAA was the only hormone with an inductive effect on carotenoid accumulation. As such, we investigated the molecular mechanism behind this response by using comparative RNA-Seq of cultures under conditions inducing maximal carotenogenesis (5 μM, day 3).

Differential expression of genes in the carotenoid pathway was determined *in silico* and verified via quantitative reverse transcriptase PCR (qRT-PCR) in treated vs. untreated cells. Transcriptome-wide changes brought about by IAA treatment were investigated using the RNA-Seq data and auxin-related genes were identified through homology matching to online databases. Assembled sequences were made available in the NCBI database, allowing future studies to use our data as a discovery platform. To our knowledge, no transcriptome-wide information currently exists in the literature regarding auxin-responsive genes in microalgae.

## Material and Methods

### Algae cultivation and strain information

*Chlorella* sp. BR2 (Genbank: JQ423151) was originally isolated from the Brisbane River (27°31’21.36”S, 153°0’32.87”E; Lim et al., 2012) and has since been used commercially. Protocols for obtaining and maintaining pure cultures were described previously by Duong et al. (2012). *Chlorella* sp. BR2, cultures were inoculated from master cultures in 250 mL flasks with BBM medium (Nichols 1973; Nichols and Bold 1965). Fluorescent lights were used a source of illumination with a light intensity of 120 μmol photons m^−2^ s^−1^ on 16-8 h light-dark cycle at 25°C with constant aeration. Phosphate and nitrate levels were determined every alternative day (using API nutrient kit following the manufacturer’s protocols) to ensure the cultures were not deprived of these essential factors. Culture medium pH was frequently measured by a pH meter (pHMaster BIO, Dynamica Scientific Ltd., UK). The culture’s optical densities were measured at 440 nm by a UV-Vis (Hitachi U2800) spectrophotometer (1 mg dry weight (DW)/mL culture is approximately equal to an OD of 2.5).

### Sampling schedule/ hormone treatments

*Chlorella* sp. BR2 cultures were grown separately (n=3 per treatment). During the exponential phase, cultures (were treated with 5, 10, 20, 50 and 100 of μmol/L IAA; 20, 100, 500 and 100 μmol/L of ABA; 50, 100, 500 and 1000 μmol/L of SA, and 10, 100, 500 and 100 μmol/L of MJ. IAA, ABA and MJ were dissolved in 100% ethanol, while SA was dissolved in deionized water. The same amounts of 100 % ethanol and deionized water were added to separate cultures (n=3) and grown as controls. Media was added regularly (addition of 25 mL of 10x BBM stock solution to 250 mL culture) to prevent any nutrient starvation effects on carotenoid accumulation. The biomass collection was performed at an interval of 2 days after the treatment by hormones and included harvesting by centrifugation (3000 ×; g; at 25°C for 3 min) and washing in Milli-Q water. The harvested biomass was freeze-dried and then stored at −20°C until carotenoid extraction took place.

### Analyses of carotenoids

Carotenoids were extracted and analyzed as described by Ahmed et al. (2014). Briefly, a mortar and pestle were used to crush the freeze-dried biomass, which was then weighed (10-20 mg). A total of 10 mL acetone was added to the biomass followed by vortexing. The, 5 mL NaCl (10% w/v), and 10 mL hexane were added, followed by vortexing and centrifugation (3000 ×; g at 4°C for 5 min). Supernatants were removed and dried in a centrifugal evaporator. A volume of 2.5 mL methanol/dichloromethane (50:50, v/v) was added to the dried supernatants for High Performance Liquid Chromatography (HPLC) analysis. The following gradient was used to separate the individual carotenoid compounds: 0 min, 80 % phase A; 48 min, 20 % phase A; 49 min, 80 % phase A; and 54 min, 80 % phase A (phase A, 92 % methanol/8 % 10 mM ammonium acetate; phase B, 100 % methyl tert butyl ether). A volume of 20 μL was injected onto a YMC C30 Carotenoid Column (3 μm, 4.6×250 mm; Waters, Milford, MA, USA). Detection was carried out using a photodiode array detector. Furthermore, a mass spectroscopy (MS) scan was undertaken using an Acquity UPLC H-Class system connected to a Quattro Premier triple quad (Micromass MS Technologies, Waters Corporation, Milford, MA, USA). Carotenoid identification was carried out by comparison of retention times, UV/Vis spectra and mass spectra against authentic standards. Identified carotenoids were quantified using individual calibration curves and concentrations were expressed as mg carotenoid per g DW.

Sampling of RNA was conducted on same-day controls and samples from the time point with the highest significant differences in total carotenoids (5 μM, day 3). A sample of 10 mL of culture was collected by centrifugation (10,000 x g, at 25°C for 7 min) from each replicate, the supernatant discarded and collected cell pellets immediately flash-frozen with liquid nitrogen and stored at −80°C. Prior to RNA extraction, cell pellets were resuspended in lysis buffer (SV Total RNA Isolation System, Promega) and then ground using a micro pestle. Total RNA was then extracted following the manufacturer’s instructions, with the exception that the incubation at 70°C was carried out at room temperature. Total RNA was not pooled but kept as respective replicates and then stored at −80°C. These RNA samples were then submitted to AGRF (Melbourne, VIC, Australia) for sequencing. All 100 bp single end reads were generated on an Illumina Hi-seq 2500 using the TruSeq3 stranded protocol. Six FastQ files were produced using Illumina control protocol version 2.2. These raw reads were then deposited in the NCBI database in order to make them available to other researchers (BioProject: PRJNA389111).

### Read quality control and trimming

The six raw read data files from sequencing were assessed using FastQC, v0.11.5 (available from: http://www.bioinformatics.babraham.ac.uk/projects/fastqc/). Sequence bias was observed in the first 13 nt at the 5’ end of the reads. This phenomenon has been described in a previous analysis of Illumina hexamer priming, which is utilized in TruSeq3 (Hansen et al. 2010). The authors recommend a correction scheme that involves reweighing transcript abundance estimates, referencing a ‘base truth’ sequence k-mer content derived from real world data. As the kallisto transcript quantification method relies upon k-mer mapping and counting to calculate expression values, these 13 nucleotides were omitted from the 5’ end of all reads prior to treatment with Trimmomatic v0.36 (Bolger et al. 2014). Trimmomatic was applied with the settings (MINLEN: 25, LEADING: 5 and SLIDINGWINDOW 4:5). This was in accordance with both the tool’s manuscript and a systematic review of Trimmomatic settings and their impact upon the quality of *de novo* assemblies of Illumina RNA-Seq data (MacManes 2014).

### Transcriptome assembly and annotation

Six FastQ formatted input read files per species were pooled and assembled in Trinity (v 2.3.2) using the following non-default settings (min_kmer_cov 2) on the UQ Flashlite HPC (Grabherr et al. 2011). Transcriptome assembly statistics were computed with the TransRate tool in assembly mode (Smith-Unna et al. 2016).

Assembled transcripts were then annotated using the Trinotate pipeline. Transdecoder (v3.0.0) was used to predict the highest-scoring open reading frames (ORFs) for each transcript with default settings as described in the Trinity supplementary material (Haas et al. 2013). The protein and transcript sequences were compared to all taxonomic groups except Metazoa (tax id: 33208) in the Uniprot Swiss-Prot database, and to *Chlorophyta* (taxid: 3041) in the TrEMBL database using BLASTp and BLASTx (Feb 28, 2018; e < 1^−5,^) (Apweiler et al. 2004). Gene Ontology (GO), eggNOG and KEGG Orthology (KO) terms were assigned to proteins based on their BLASTp best hits (Apweiler et al. 2004; Ashburner et al. 2000; Powell et al. 2011). Hmmer (v3.1b2) was used to find protein domain matches with the Pfam database (Mar 8, 2018; DNC = 10^−5^) (Finn et al. 2013; Finn et al. 2011). Proteins were also scanned for putative signal peptides and transmembrane domains with tmHMM (v2.0c) (Krogh et al. 2001). Finally, rRNA contamination was investigated using RNAmmer (v1.2) (Lagesen et al. 2007). These results were then compiled into a CSV file for use in subsequent analyses which is available from http://schenklab.com/transcriptomic_analysis/chlorella_br2/

### Transcript quantification and differential expression analysis

An overview of the functional enrichment analysis is provided in Figure 1. First, transcripts were quantified using the kallisto tool (Bray et al. 2016). Kallisto uses a novel k-mer-based approach termed ‘pseudoalignment’ to quantify assembled transcripts against a library of differentially expressed k-mers generated from RNA-Seq reads. This method is more accurate than traditional computationally expensive alignment algorithms employed in TopHat-Cufflinks or Bowtie-RSEM workflows, which function by mapping all of the read input data to the assembled transcriptome or a reference and finding regions with differentially expressed features. In all instances, trimmed high quality reads were used as the input dataset, following the recommendations of the kallisto documentation. DEseq2 was used to infer differential expression of the kallisto-computed TPM (transcripts per million) using the Bioconductor package for R (Love et al. 2014).

**Figure 1.**
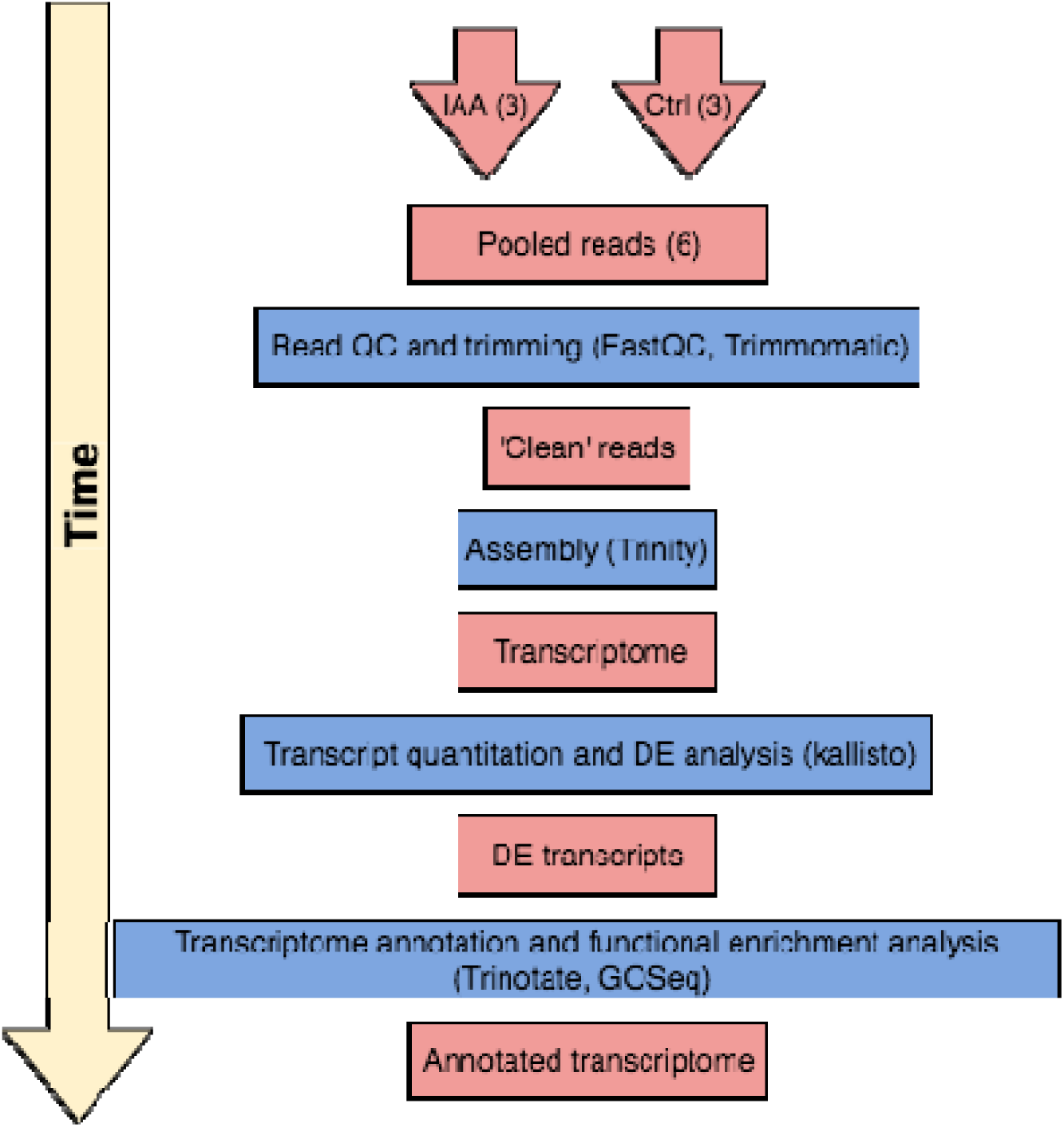
Assembly and functional enrichment analysis pipeline of RNA-Seq data obtained from 3 IAA-treated and 3 untreated *Chlorella* sp. BR2 cultures.

GO terms were examined for functional enrichment using the GOseq Bioconductor package to independently compare the significantly up- and down-regulated gene sets to the assembly. The GO enrichment and depletion results were then submitted to the CateGOrizer web tool for classification into Plant GOSlim lexicon (Available from: http://www.animalgenome.org/tools/catego/).

### qRT-PCR confirmation of differentially expressed genes

To confirm the differential expression calls generated *in silico*, a series of 22 primer sets were designed within the alignment region of their respective transcript to the Swiss-Prot database using Primer BLAST (Ye et al. 2012). These primer sequences and information regarding their targets is available in Supplementary Table 1. RNA was extracted from independently treated microalgal samples using the Promega Maxwell^®^ Plant RNA extractor as per the manufacturer’s instructions. Extracted RNA was quantified and qualitychecked on a NanoDrop™ One spectrophotometer to ensure samples were of sufficient quality for subsequent analyses. cDNA was synthesized using Superscript III (Invitrogen). qRT-PCR was carried out using a 10 μL reaction volume that contained cDNA generated from 10 ng RNA, 5 μL SYBR Green and 1 μL 100 nmol primers (each). Melt curve analysis and agarose gel electrophoresis were used to determine whether transcript amplification was successful. To quantify relative gene expression, the individual efficiency corrected calculation delta Ct method was used with *GAPDH* as the reference gene (Rao, Huang et al. 2013).

### Statistical analyses

For carotenoid measurements, mean ± standard SEs were derived for all data and were statistically analysed with two-way ANOVA (GraphPad Prism 7.03). Tukey’s HSD test was used to test the differences among groups of different trials. Values of *p* < 0.05 were considered to be statistically significant. For qRT-PCR quantifications, a Student’s t-test was performed on the mean relative expression ratio to the housekeeping gene, and *p* values < 0.05 were considered statistically significant. For RNA-Seq quantitations a False Discovery Rate (FDR) < 0.05 was considered significant.

## Results

### IAA effect on growth and carotenoid accumulation in *Chlorella* sp. BR2

The plant hormones IAA, SA, ABA, and MJ were applied to selected microalgal cultures to examine potential inductive effects on growth and carotenoid content. Growth of *Chlorella* sp. BR2 was unaffected by application of IAA, as the optical densities of control cultures ranged from 1.15 ± 0.01 on day 1 to 1.58 ± 0.3 on day 5, while the IAA-treated cultures were in a similar range (1.21 ± 0.4 on day 1 to 1.54 ± 0.02 on day 5).

Carotenoid accumulation in response to plant hormones differed in the four plant hormone treatments. The total carotenoid production increased in *Chlorella* sp. BR2 after treatment by IAA from day 1. Significantly higher total carotenoid production compared to the control group was observed on day 3 at 5, 50 and 100 μmol/L and day 5 at 5 μmol/L (2.65 ± 0.14, 2.71 ± 0.01 and 2.87 ± 0.13 vs. 1.11 ± 0.16 mg/g DW in control cultures for day 3; 3.07±0.62 vs. 0.88 ± 0.18 mg/g DW in control cultures for day 5; Tukey’s test, *p* < 0.05; Figure 2). However, no significant increase in total carotenoid production was found up to 5 days after treatment by MJ, SA and ABA in *Chlorella* sp. BR2 (two-way ANOVA, *p <* 0.05; Figure 2).

**Figure 2.**
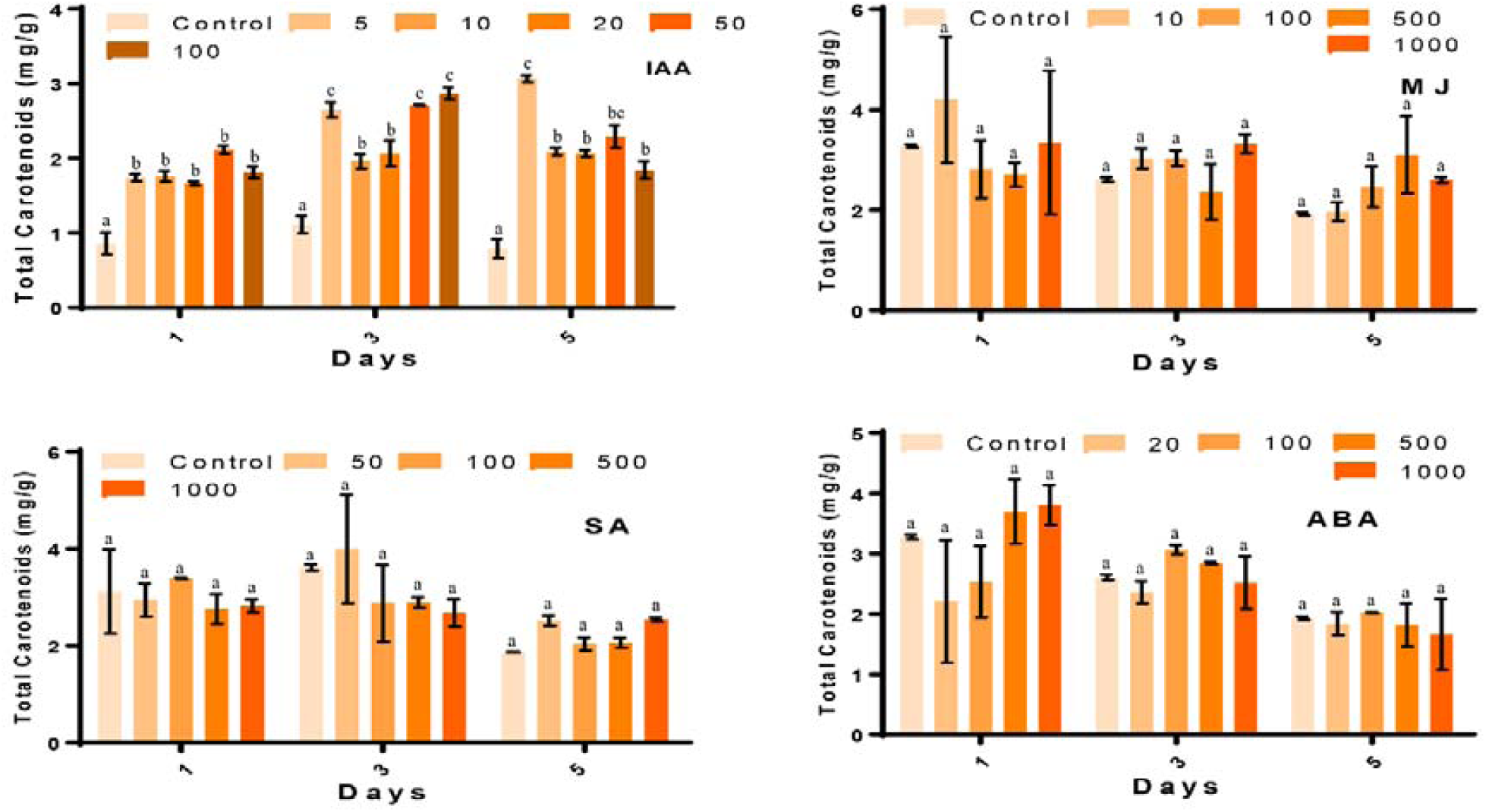
Effect of different concentrations of IAA, MJ, SA and ABA on total carotenoids in *Chlorella* sp. BR2. Values are the mean ± SEs from three separately grown cultures (ANOVA test; *p* < 0.05). Values with different letters are significantly different.

HPLC analysis showed three major carotenoids, violaxanthin, lutein and β-carotene, in *Chlorella* sp. BR2 treated with IAA. Lutein was the major carotenoid quantified in all treated and untreated samples. When compared to controls, the highest content of violaxanthin was found on all test days with 100 μmol/L IAA (day 1:0.89 ± 0.01 vs. 0.25 ± 0.14 mg/g DW; day 3: 0.89 ± 0.01 vs. 0.32 ± 0.04 mg/g DW; day 5: 0.92 ± 0.04 vs. 0.32 ± 0.04 mg/g DW) (Tukey’s test, *p* < 0.05; Figure 3). The highest lutein contents were found on day 3 (5 μmol/L: 1.4 ± 0.04; 50 μmol/L: 1.5 ± 0.01; 100 μmol/L: 1.72 ± 0.06 vs. control 0.56 ± 0.1 mg/g DW), and day 5 (5 μmol/L: 1.73 ± 0.02 vs. control 0.31 ± 0.17 mg/g DW) (Tukey’s test, *p* < 0.05; Figure 3). The highest β-carotene content was found at day 3 at 5 μmol/L (0.69 ± 0.11 vs. control 0.23 ± 0.02 mg/g DW) (Tukey’s test, *p* < 0.05; Figure 3).

**Figure 3.**
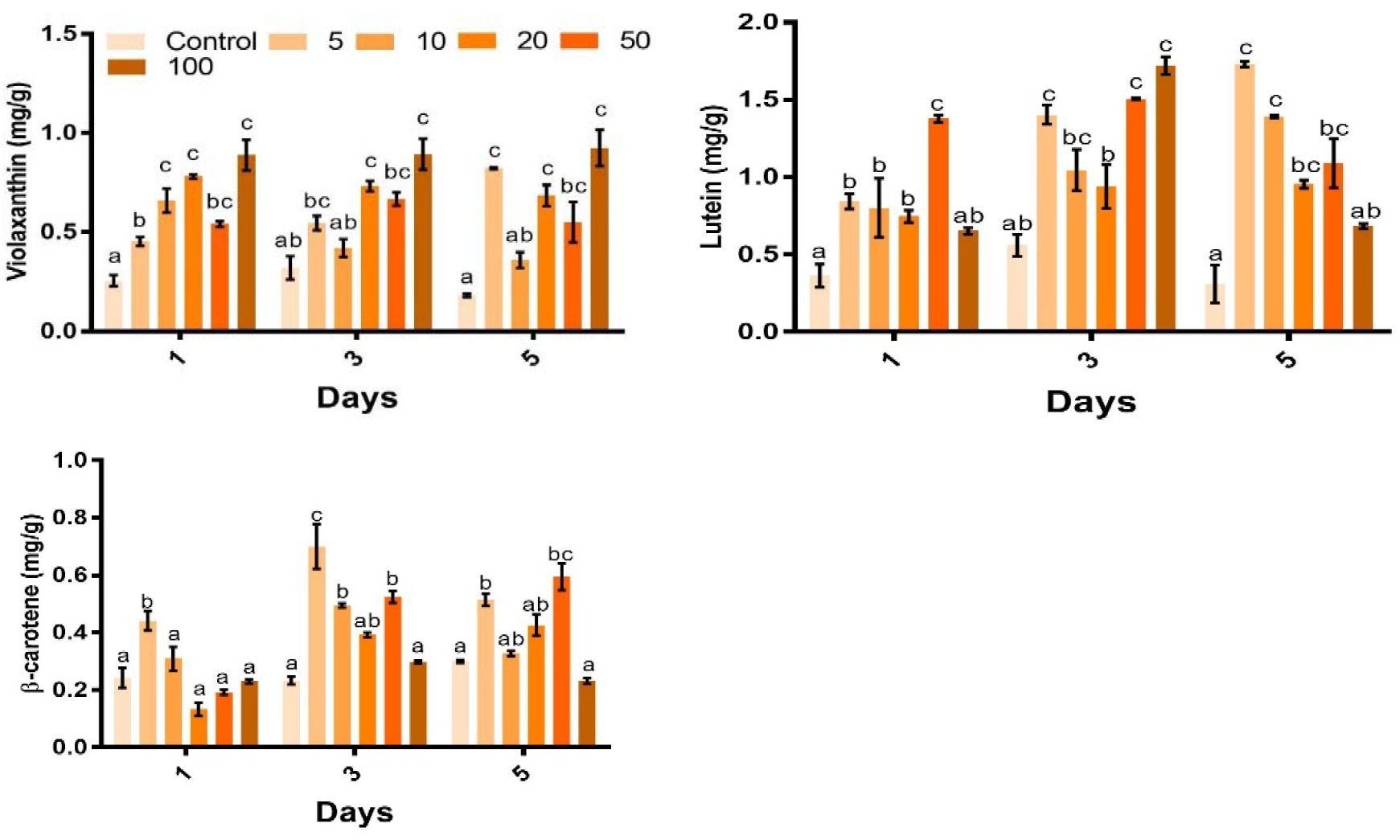
Effect of different concentrations (μmol/L) of IAA on violaxanthin, lutein and β-carotene contents of *Chlorella* sp. BR2. Shown are mean values ± SEs from three separately grown cultures. Values with different letters are significantly different.

### Transcriptome assembly and quality assessment

Pooling the trimmed data from all six samples yielded 11.2 Gbp of high quality sequence from 129 million reads. This was assembled into an 80 Mbp transcriptome consisting of 113,167 transcripts and 92,024 genes, with an N50 value of 1,036 bp and average coverage of 140-fold (Table 1). Although N50 is a common statistic for comparing transcriptome assemblies, it does not take into account factors such as transcriptome completeness. In order to assess transcriptome completeness, Benchmarking Using Single Copy Orthologues (BUSCO) v2.0 analysis was used with the Eukaryota (May, 2017) test set (Simão et al. 2015). Bowtie2 was also used with default settings to map trimmed reads to the assembly and determine map-back rate (97.4%) (Langmead and Salzberg 2012). The BUSCO results shown in Figure 4, when taken in conjunction with the high coverage depth and map-back percentage, suggest that the obtained sequencing data captured a wide representation of the expressed transcriptome.

**Table 1.** Read and assembly summary statistics for *de novo* assembly of *Chlorella* sp. BR2.

| Sample | 1 | 2 | 3 | 4 | 5 | 6 |
| --- | --- | --- | --- | --- | --- | --- |
| BioSample | SAMN07280588 | SAMN07280589 | SAMN07280590 | SAMN07280591 | SAMN07280592 | SAMN07280593 |
| Treatment | Control |  |  | IAA |  |  |
| Reads Before | 22,873,627 | 22,041,700 | 21,217,208 | 20,083,170 | 19,922,057 | 22,735,553 |
| Total | 66,132,535 |  |  | 62,740,780 |  |  |
| Reads After | 22,869,528 | 22,033,781 | 21,212,149 | 20,077,616 | 19,917,129 | 22,730,709 |
| Total | 66,115,458 |  |  | 62,725,454 |  |  |
| Nuc Before | 2,287,362,700 | 2,204,170,000 | 2,121,702,800 | 2,008,317,000 | 1,992,205,700 | 2,273,555,300 |
| Total | 6,613,235,500 |  |  | 6,274,078,000 |  |  |
| Nuc After | 1,984,523,686 | 1,912,112,197 | 1,840,773,015 | 1,742,422,858 | 1,728,305,346 | 1,973,271,103 |
| Total | 5,737,408,898 |  |  | 5,443,999,307 |  |  |
|  | 11,181,408,205 |  |  |  |  |  |
| Assembled Nucleotides (%) | 80,027,350 |  |  |  |  |  |
| Trinity genes | 92,024 |  |  |  |  |  |
| Trinity transcripts | 113,167 |  |  |  |  |  |
| N50 | 1,036 |  |  |  |  |  |
| Mean Tx length | 707 |  |  |  |  |  |
| ORFs | 73,809 |  |  |  |  |  |
| Complete ORFs | 13, 112 (18 %) |  |  |  |  |  |
| Estimated coverage | 140 |  |  |  |  |  |
| Read map-back percentage | 97.43% |  |  |  |  |  |

**Figure 4.**
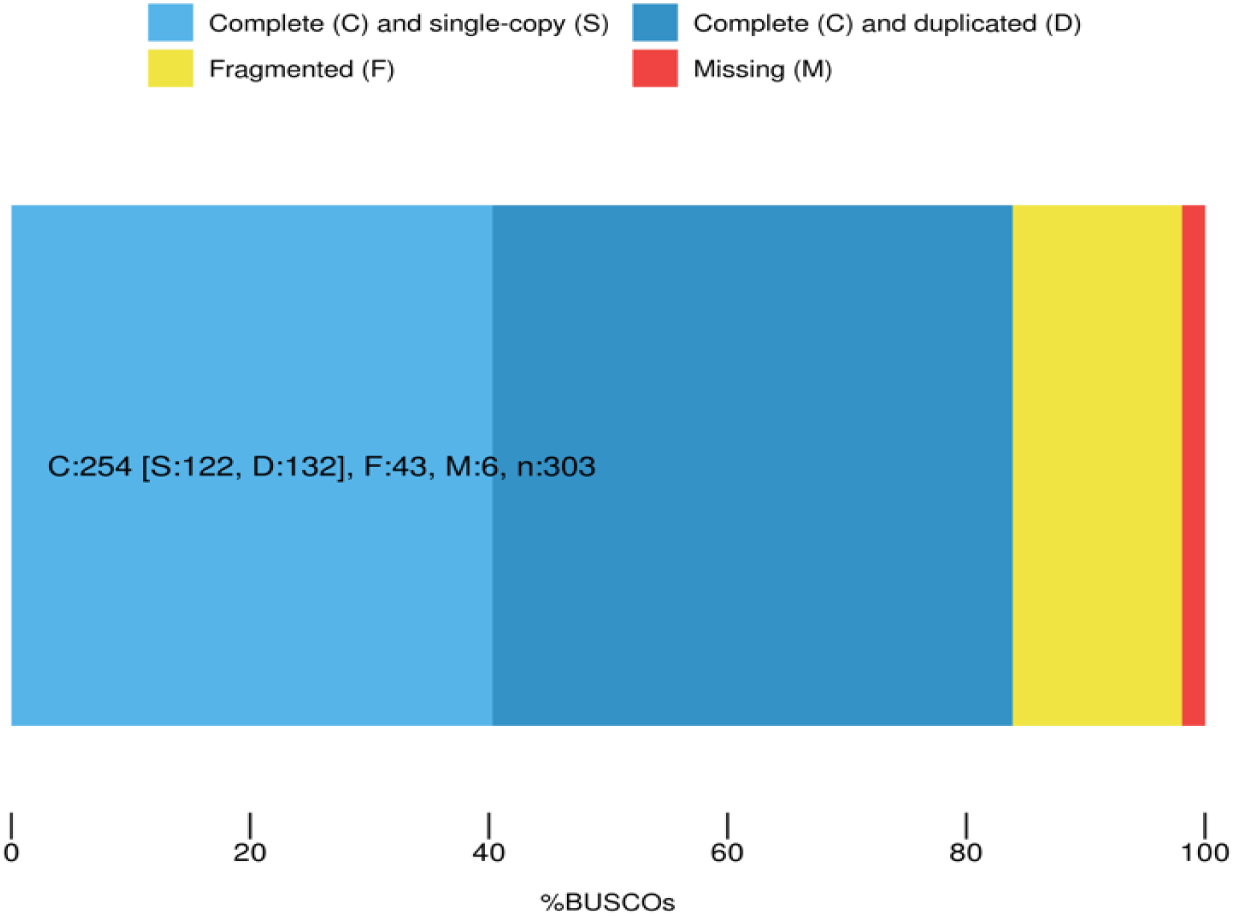
BUSCO analysis of conversed core genes in assembled *Chlorella* sp. BR2 transcriptome.

### Transcript annotation and differential expression analysis

Of the assembled transcripts, 68,478 (60 %) were predicted to encode 73,809 ORFs, of which 13,112 (18 %) were full length (Table 1). A total of 56,466 transcripts (50 %) were assigned either descriptions (BLAST), conserved protein domains (Pfam) or functional annotations (BLASTx). A detailed description of these annotations can be found in Figures 5 and 6. Of the 16,404 transcripts considered to be differentially expressed (15 % of the assembly), 4,189 (6 %) were significantly downregulated, while 3,178 (5 %) were significantly upregulated as determined using DESeq2 (FDR < 0.05). This subset of genes was considered as the differentially expressed set in subsequent functional enrichment analyses.

**Figure 5.**
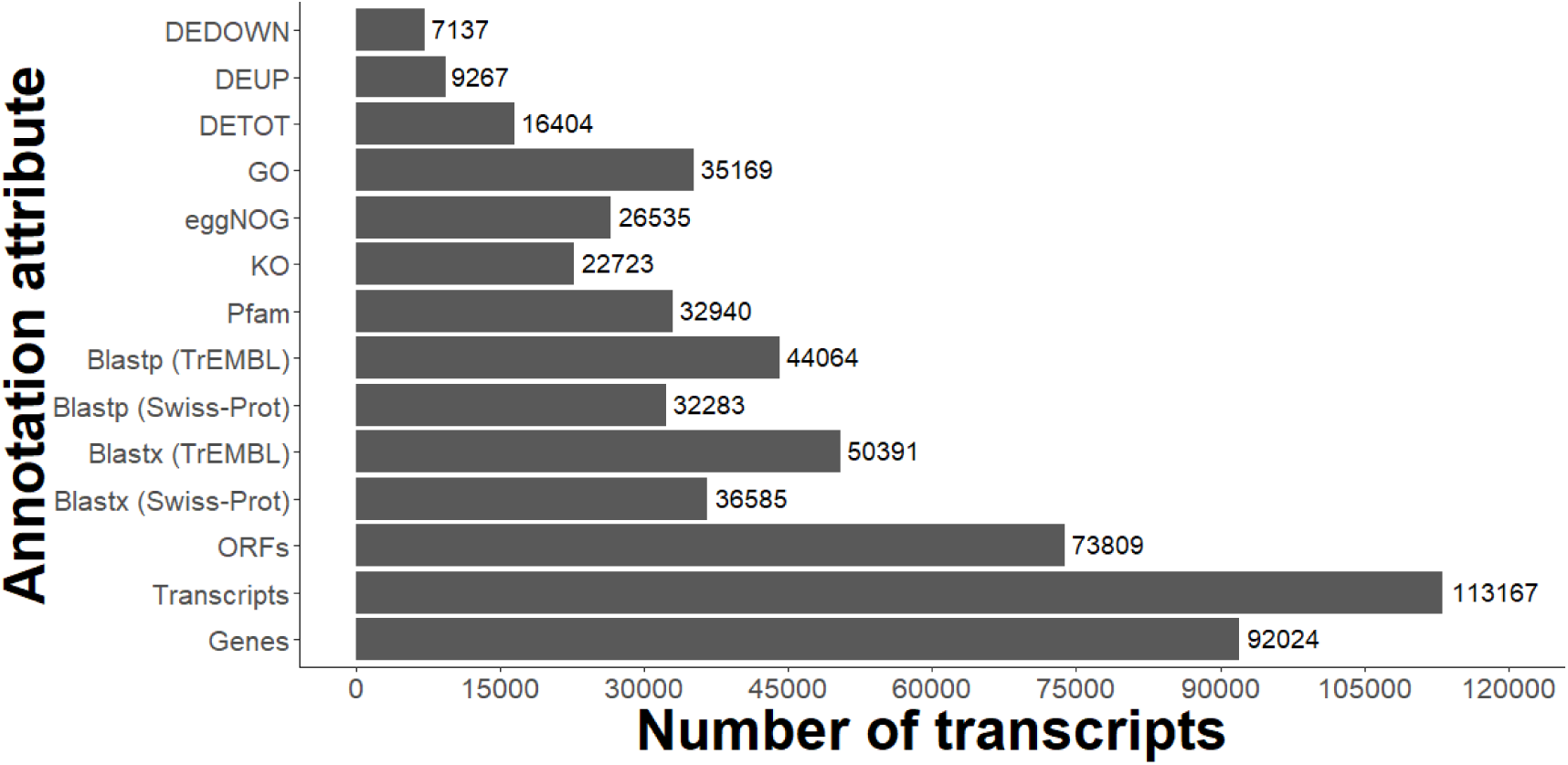
Summary of annotation attributes allocated to transcripts of the *Chlorella* sp. BR2 transcriptome by the Trinotate pipeline.

**Figure 6.**
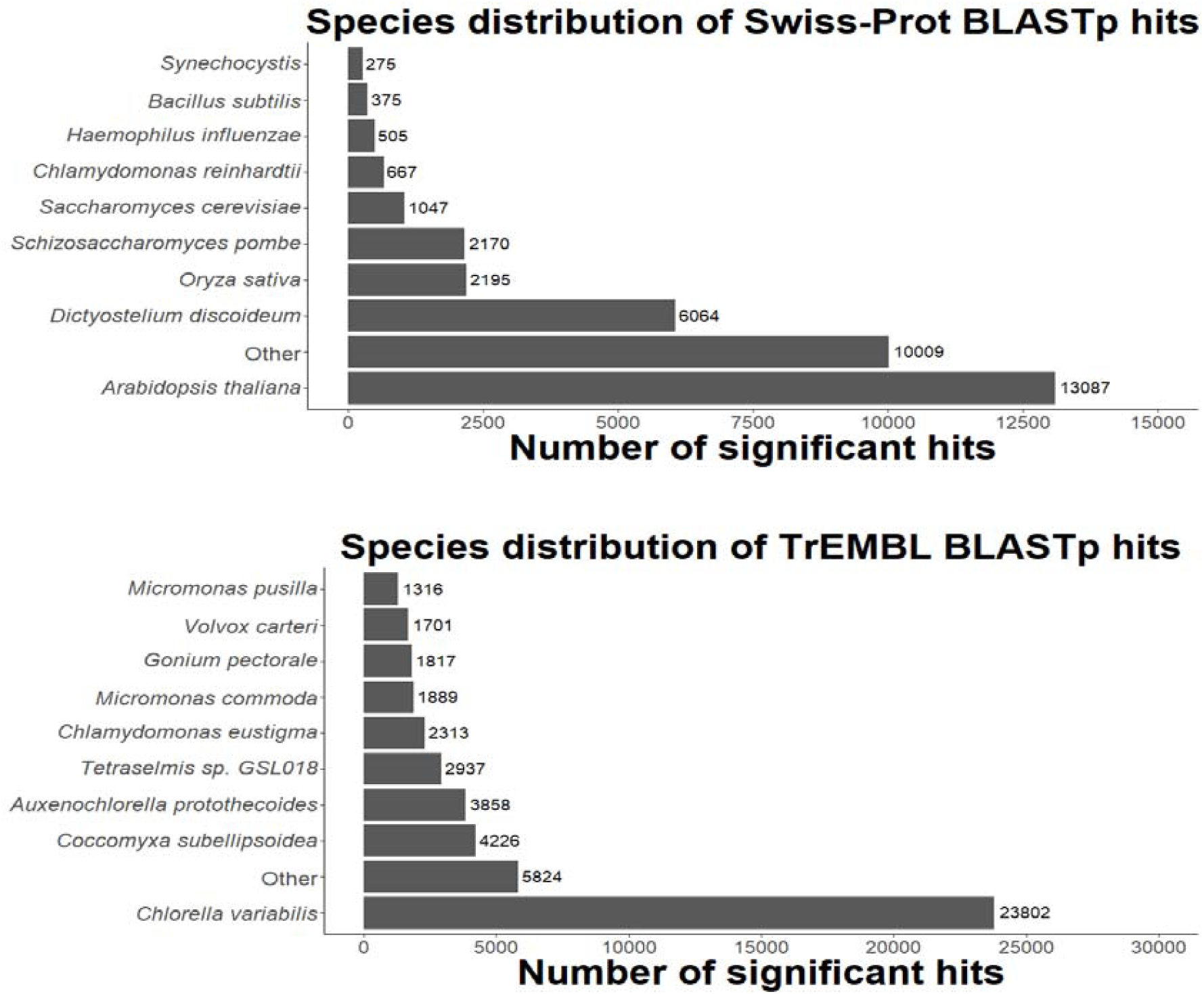
Breakdown of BLASTp matches to the Swiss-Prot and TrEMBL databases by species of the matched sequence.

### GO enrichment analysis

The GOSeq Bioconductor package for R was used to independently test the up- and down-regulated transcript sets for enriched GO annotations (FDR < 0.05). In the upregulated transcripts, 186 GO terms were enriched and assigned to 103 Plant GOslim terms (Supplementary Figure 1). The most frequent terms above the root level included ‘biosynthetic process’, ‘nucleobase process’ and ‘cellular component organisation and biogenesis’. Of interest to this study were ontologies such as ‘lipid metabolic process’ (14), ‘response to stress’ and ‘signal transduction’ (12). In addition, 924 GO terms were enriched in the downregulated transcripts and assigned to 103 Plant GOSlim terms. The most frequent annotations above the root level were ‘cellular component organisation and biogenesis’ (87), ‘binding’ (74) and ‘nucleobase process’ (66). ‘signal transduction’ (32), ‘response to stress’ (25), ‘lipid metabolic process’ (18) also featured in this dataset (Supplementary Figure 2).

### Identification and expression of auxin-inducible transcription factors

Assembled transcriptomes were submitted to the PlantTFDB v4.0 online tool for transcription factor (TF) prediction (Jin et al. 2017). 528 transcripts were found to encode proteins showing homology to known algal TFs. Of these transcripts, 37 (7 %) and 48 (9 %) were up- and down-regulated, respectively. Figure 7 shows the distribution of the differentially expressed TFs. The most prevalent TF family was *GATA*, followed by *bZIP, C3H, MYB* and *ERF*.

**Figure 7.**
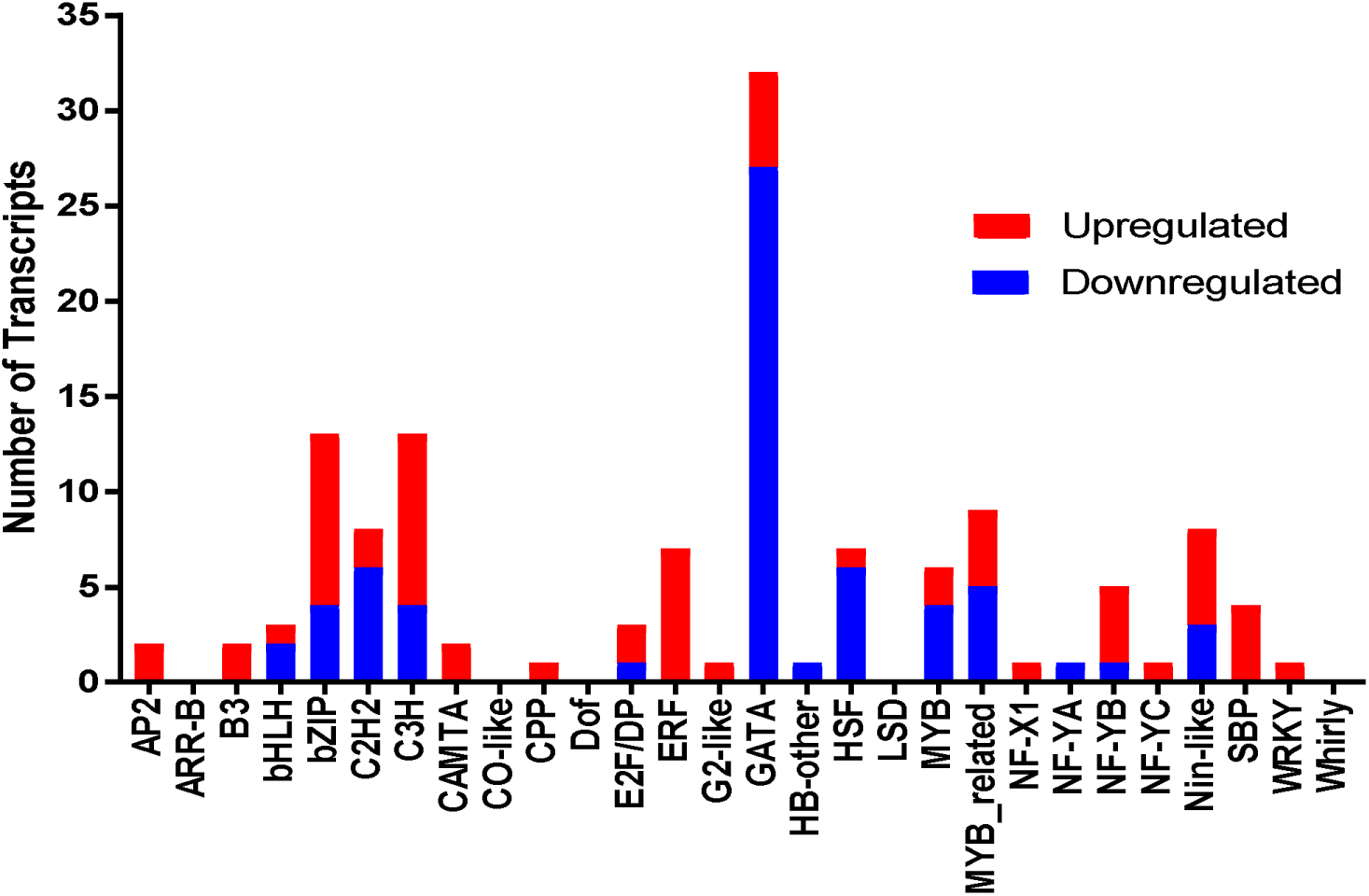
Distribution of differentially expressed transcription factors under auxin treatment in *Chlorella* sp. BR2.

### Identification and differential expression of carotenogenic transcripts

A total of 109 protein-coding transcripts were assigned to 27 genes associated with the MEP (non-mevalonate) or carotenoid biosynthetic and catabolism pathways based on their BLAST matches (Supplementary Table 2). Of these transcripts, 22 (20 %), belonging to 12 (41 %) of the genes displayed differential expression under auxin treatment. 19 (86 %) transcripts were considered as upregulated, while 4 (18%) were downregulated. An overview of the reconstructed carotenoid pathways is shown in Figure 8.

**Figure 8.**
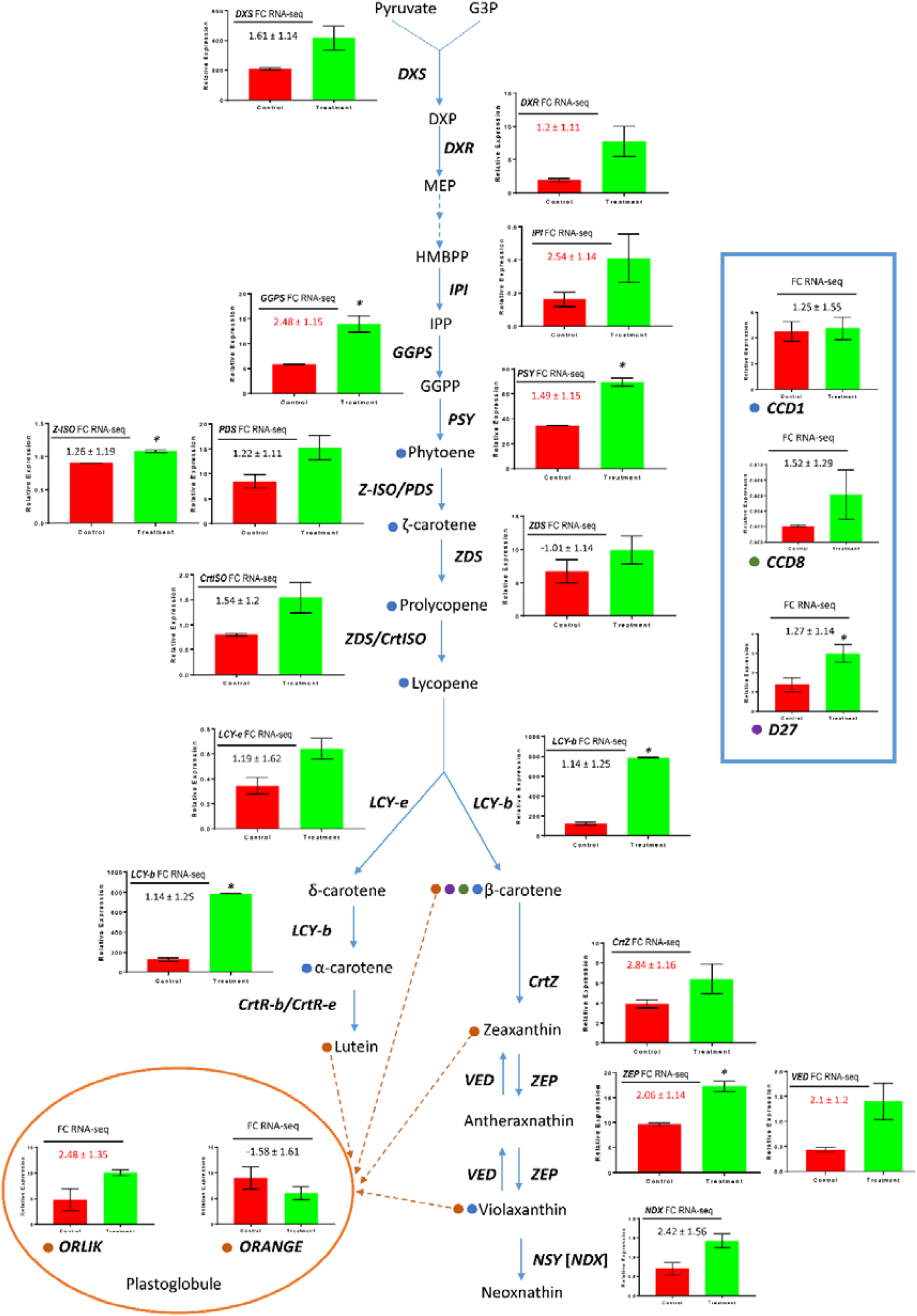
Pathway diagram and gene expression of reconstructed carotenoid-related processes, including biosynthesis, degradation and storage for *Chlorella* sp. BR2 under IAA treatment. Inserted graphs show qRT-PCR analyses of the RNA used for RNA-Seq, with asterisks (*) indicating significant differences (Student’s t-test; *p* < 0.05). The inserted tables show RNA-Seq expression data for each gene, with red numbers indicating significant differences (FDR < 0.05). Colored circles indicate association between a given carotenoid-gene pair.

### qRT-PCR confirmation of differentially expressed transcripts

qRT-PCR was used to confirm the existence and measure the relative expression of transcripts belonging to carotenogenic pathways. All PCR reactions produced a single amplicon, as assessed by melt-curve analysis and agarose gel electrophoresis. Statistically significant upregulation was observed in six transcripts (*GGPS, PSY, Z-ISO, LCYb, ZEP* and *D27*), which were upregulated 2.38, 2.01, 1.22, 6.29, 1.79 and 2.18-fold, respectively (Figure 8). *GGPS, PSY* and *ZEP* expression followed the same trends in both RNA-Seq and qRT-PCR data, while *Z-ISO, LCYb* and *D27* expression were not significantly differential in the RNA-Seq data. Despite the lack of statistical significance, qRT-PCR results showed that 19 transcripts followed the same trend as the RNA-Seq data, except for *ZDS*.

## Discussion

In this study, next generation RNA sequencing was used to generate a reference transcriptome for *Chlorella* sp. BR2 and to characterize transcriptional changes in response to IAA treatment. Notably, this study is the first published transcriptome assembly of an Australian microalgal species, apart from the marine diatom *Ceratoneis closterium*, which was performed using lower-throughput 454 pyrosequencing technology (Hook et al. 2014). Average transcript coverage in this experiment was 140-fold, which was likely responsible for the high quality scores following assembly. Several quality assessments indicated that the reference transcriptome was of high quality. As such, the assembled sequences have been made available in the NCBI database to benefit the research community. qRT-PCR results matched the sequencing data in all of the cases except for *ZDS*, although all monitored transcripts displayed general trends in line with those predicted by RNA-Seq. As qRT-PCR relies on the expression levels of the targeted amplicon being an adequate proxy for whole transcript expression, it may be possible that the discrepancy between qRT-PCR and RNA-Seq data is due to primer sequences targeting transcripts that are members of gene families. RNA-Seq quantitations do not suffer from this issue, as kallisto only uses unique regions of transcript-read pseudoalignments to feed its statistical model. Off target amplification could be investigated by designing primer sets that span the entire length of transcripts of interest and then comparing relative expression levels.

### IAA-induced carotenoid profiles and associated gene expression

All IAA treatments (5 – 100 μmol) promoted increases in carotenoid content on days 1, 3 and 5. The profiles of carotenoids became more diverse in a dose-dependent manner; the relative amount of violaxanthin increased with dose, while lutein decreased relative to violaxanthin, except for 5 μmol IAA, where both were present at similar levels, but higher than the control cultures on days 3 and 5. No difference was found across days in control samples, suggesting that the modification of carotenoid profile was due to the action of IAA and not culture conditions. These results are in agreement with a previous study of *Chlorella vulgaris* that observed a 150 % increase in total carotenoid content 48 h following treatment with 0.1 μmol IAA (Piotrowska-Niczyporuk and Bajguz 2014). No carotenoid profiling was conducted in their study, which prevented comparisons to this aspect of our data being made.

In order to understand the observed IAA-responsive carotenoid accumulation at the transcript level, we examined the relative expression of genes in the carotenoid biosynthetic pathway. Expression of several rate-limiting transcripts in the MEP and carotenoid pathways were upregulated upon IAA treatment, including *DXS, DXR, IPI* and *GGPS*. In the MEP pathway, *DXS* catalyzes the first step by condensing a molecule of pyruvate and glyceraldehyde 3-phosphate to form 1-deoxy-D-xylulose 5-phosphate (DOXP), which is then reduced by *DXR* to 2-C-methylerythritol 4-phosphate (MEP). The final step of this pathway is also rate-limiting, involving the isomerization of isopentenyl pyrophosphate (IPP) to dimethylallyl pyrophosphate (DMAPP) by *IPI*. Rate-limiting enzymes of the carotenoid biosynthesis pathway include *GGPS, PSY, PDS* and *ZDS*. *GGPS* produces geranylgeranyl pyrophosphate (C_20_) from 3 molecules of IPP (C_5_) and 1 DMAPP (C_5_). While geranylgeranyl pyrophosphate serves as a precursor for many terpenes, *PSY* represents the first committed step of the carotenoid biosynthetic pathway, where it forms phytoene (C_40_) from 2 geranylgeranyl pyrophosphates. Phytoene is then desaturated in two steps by *PDS* to ζ-carotene and then to lycopene in another two steps by *ZDS*. Lycopene is a branching point in the pathway, where the relative amounts of *LCYb* and *LCYe* determine the ratio of α-carotenoids (lutein) to β-carotenoids (β-carotene) and xanthophylls (zeaxanthin, violaxanthin and neoxanthin). We found that levels of *GGPS, PSY* and *PDS* transcripts increased at day 3 for the treatment group. This pattern of expression is consistent with those observed under several conditions that promote carotenoid accumulation in microalgae, such as nitrogen deprivation and high light intensity in *Haematococcus pluvialis* (Ma et al. 2017). Our data also showed that *LCYb* and *LCYe* both increased in expression, although the induction was more pronounced in *LCYb*. This was observed concomitantly with an increase in total carotenoid and the relative proportion of lutein, at the expense of violaxanthin. When taken together, these results suggest that *LCYb* likely determines the total carotenoid level in *Chlorella* sp. BR2 through its broad action on both branches of the pathway, while *LCYe* regulates the relative α-carotenoid level by directing carbon flux through the α-carotene branch of the pathway through its specific action at the beginning of this section. This is consistent with a study in maize endosperm, which found that the ratio of *LCYb* to *LCYe* mRNA was a determinant of α-to-β carotene ratio (Naqvi et al. 2011). In contrast with other physical and chemical stressors such as high irradiance, nitrogen deprivation and high temperature, IAA-induced carotenoid accumulation occurred without attenuated growth of *Chlorella* sp. BR2. This observation has tremendous implications for algal culture practices; if mutants hypersensitive to endogenous auxin could be isolated, it may be possible to genetically engineer a strain of microalgae containing much higher levels of carotenoids and very long chain-polyunsaturated fatty acids (VLC-PUFAs), without sacrificing growth characteristics under inductive conditions.

### Cross-talk of carotongenesis with phytohormone signaling and other pathways

Most photosynthetic organisms feature mechanisms to negate the biological impact of excess irradiance, which is the production of reactive oxygen species (ROS). This chiefly occurs through non-photochemical quenching (NPQ), where incidental photons are absorbed and reemitted as heat. The xanthophyll cycle is present in chlorophytes and land plants and typically features the interconversion of zeaxanthin to antheraxanthin and violaxanthin through successive epoxidation and de-epoxidation reactions (Latowski et al. 2011). The rate-limiting step of this process has been shown to be the de-epoxidation of violaxanthin, with the relative amount of *VIO* to *ZEP* expression, and consequently, the level of NPQ-active zeaxanthin, determining the amount of NPQ (Chen and Gallie 2012). qRT-PCR and RNA-Seq analysis in the present study found that both *VIO and ZEP* expression increased by similar amounts (∼2-fold; Figure 8). As this change of expression was of a similar magnitude, the results suggest that IAA application did not affect the level of NPQ, and that transcript increases were most likely due to the increase in total flux through the β-carotenoid biosynthetic pathway. Violaxanthin also serves as a substrate for the synthesis of neoxanthin by *NSY*, and is ultimately converted into the stress hormone ABA by *NCED* and *AAO* (Finkelstein 2013). Overexpression of *ZEP* has been shown to enhance formation of neoxanthin and ultimately confer tolerance to conditions such as drought and salinity in *Arabidopsis thaliana* through active ABA biosynthesis (Park et al. 2008). We also found that the transcript level of *NSY* increased at least 2-fold in the treatment groups with qRT-PCR and RNA-Seq, although this did not achieve statistical significance in the qRT-PCR samples. Therefore, it may be possible that increased carbon flux through the β-carotenoid pathway may be due to active ABA biosynthesis, rather than increased NPQ. Further experiments using pulsed amplitude modulated fluorometry and measurements of ABA will produce a direct conclusion as to whether IAA application affected the level of NPQ in *Chlorella* sp. BR2 or if increased *VIO* and *ZEP* expression was due to activation of ABA biosynthesis.

ABA has been shown to promote astaxanthin synthesis in *H. pluvialis*, through triggering transition into the stress-resistant astaxanthin producing cyst morphotype (Kobayashi et al. 1997). However, ABA itself did not have an effect on carotenoid accumulation in *Chlorella* sp. BR2 at the doses examined in this study, This is in contrast with a previous study on *Chlorella pyrenoidosa* ZF, where 0.38, 1.89 and 3.78 μmol/L doses increased the carotenoid content (1.6-fold), 37.8 and 75.6 μmol/L had no effect, and 189 μmol/L suppressed production (3-fold). The lowest concentration of ABA trialed in this study was 20 μmol/L, which may have been above the activity window for ABA-inducible carotenoid formation in *Chlorella* sp. BR2.

ABA treatment produces common phenotypes with auxin such as salt tolerance and lipid accumulation in plants and microalgae (Tarakhovskaya et al. 2007). We also observed IAA-triggered downregulation of transcripts shown to be downregulated by ABA in *Chlorella sorkinara* RNA-Seq data, such as *DIVINYL CHLOROPHYLLIDE A 8-VINYLREDUCTASE*, (Khasin et al. 2017). This supports the notion that ABA signaling acts as a downstream effector of auxin signaling, which is mediated in-part by auxin-induced carotenoid synthesis. In land plants, IAA treatment heavily induces ABA biosynthesis within hours, which also activates ABA signaling (Hansen and Grossmann 2000). Microarray transcriptome profiling of IAA and ABA treated *A. thaliana* revealed that 156 of 344 auxin responsive genes were regulated by ABA, of which 62 % were regulated in the same direction (Yang et al. 2014). This study also identified transcripts that were upregulated under IAA but downregulated in public *C. sorkinara* ABA RNA-Seq data, namely *PSY* (Khasin et al. 2017). This opposing regulation may represent the existence of crosstalk between these two signaling pathways. Indeed, differential regulation of members within gene families participating in ABA signaling in response to multiple stressors and phytohormones has been documented, with characteristic transcriptional reconfigurations associated with particular conditions and hormones (Cao et al. 2016; Chan 2012).

The fact that ABA signaling components and many other genes are differentially regulated by auxin and ABA is suggestive of a signaling mechanism whereby auxin and ABA both trigger a basal ABA response, and auxin fine-tunes this response through upstream effects, such as the production of carotenoids and regulation of auxin-specific targets. This is reasonable in light of the fundamental action of auxin, which is promoting the G2-M transition of the cell cycle, facilitating cell division and expansion (Wang and Ruan 2013). Conversely, ABA acts earlier by halting the G1-S transition and thus entry into the cell cycle (Kobayashi et al. 2016). As auxin signaling promotes ABA production but also progression of the cell cycle, a regulatory circuit which can selectively activate and inhibit discrete components of ABA signaling, such as stress tolerance, under IAA, is required. Such a system is evident in the selective activation of members of ABA signal transduction components in response to varying stressors in the grass *Brachypodium distachyon* (Cao et al. 2016). A unique set of these proteins are activated upon auxin treatment, indicating that auxin-induced ABA signaling may have distinct effects to direct treatment with IAA. This is likely achieved by mutual genetic regulation which is sensitive to the relative amplitude of each signal, similar to auxin-cytokinin homeostasis in land plants, where the auxin-cytokinin ratio is a key regulator of meristem development, among other systems (Su et al. 2011).

One avenue for this crosstalk that has been proposed is ROS signaling, which is known to interact with auxin and ABA signal transduction through their mutual transcriptional changes and impacts on MITOGEN-ACTIVATED PROTEIN KINASE signaling (Sewelam et al. 2016). ROS signaling has been implicated in responses to other adverse conditions in microalgae, including high irradiance, salinity, temperature and nitrogen deprivation (Zhu et al. 2009). Further supporting the existence of an ABA-auxin-ROS interaction network is the fact that these signals affect common transcription factors, such as members of the AP2, WRKY, SBP, bZIP and MYB families, which are associated with stress responses, through distinct cis-elements (Gu et al. 2017; Xu et al. 2017). Further studies featuring null mutants for components of putative signaling pathways will provide useful for elucidating the precise nature of this interplay between hormones in microalgae, which only possess a subset of the signal transduction machinery present in land plants.

Piotrowska-Niczyporuk and Bajguz (2014) found that enzymatic and non-enzymatic antioxidant activities increased following treatment with several auxins, as measured by lipid peroxidation, ascorbate peroxidase, glutathione peroxide, superoxide dismutase and catalase levels. This is consistent with our transcriptome annotation, which found that transcripts associated with ascorbate peroxidase, glutathione peroxidase, catalase and superoxide dismutase were upregulated (Supplementary Table 3).

### Regulatory networks involved in auxin pathways

RNA-Seq data from the present study was then used to interrogate the relative enrichment of GO classifiers in an effort to identify systems regulated by auxin. Interestingly, a larger proportion of GO classifiers associated with metabolic biosynthetic pathways were downregulated than upregulated (Supplementary Figures 1 and 2). This is consistent with the general perspective of stress responses in living organisms, where it is most likely energetically favorable to shut down extraneous systems and only produce stress-related proteins. This behavior is thought to also inhibit toxic protein aggregate formation (Katz and Orellana 2012). Indeed, we found that the total number of differentially expressed transcripts (4189 vs. 3178) and GO terms (924 vs. 186) were significantly higher in the downregulated subset of transcripts when compared to the upregulated subset.

The observed variation in expression of orthologues of many auxin-inducible TFs in land plants is suggestive of a receptor-mediated, transcriptional configuration, rather than a reaction to IAA on a chemical basis. Among the auxin-responsive TFs observed in our transcriptome data, a correlation was observed with auxin-responsive TFs in land plants. For example, in this study we found that a number of AP2/ERF, bZIP, CAMTA and WRKY factors were upregulated. AP2/ERF transcriptional regulators show responsiveness to multiple plant hormones, including auxin, and were enriched in rice crown roots following auxin exposure (Kitomi et al. 2011). bZIP factors have been implicated in downstream fine-tuning of auxin-responsive gene transcription in relation to available energy through chromatin remodeling and binding to promoter elements (Weiste et al. 2017). CAMTA and WRKY mutants were shown to have lower levels of auxin-inducible transcripts in *Arabidopsis* and auxin-inducible properties in wild-type plants (Bakshi and Oelmüller 2014; Galon et al. 2010).

A common theme of the upregulated TFs in this study is that most are involved in auxin signal transduction and responses to abiotic stress. Interestingly, a large proportion of these induced transcription factors are regulated by Auxin Response Factors (ARFs) *in planta*, however our data and De Smet and Beeckman (2011) did not support the existence of ARFs, suggesting that a functionally equivalent system of unknown sequence may exist in microalgae. ARFs are known to confer their action through binding at their C-terminal region to auxin responsive elements (AuxRE), which are comprised of variations of a core ‘TGTCT’ motif, which is repeated several times in the promoter region of auxin responsive genes. De Smet and Beeckman (2011) used the lack of detectable green fluorescent protein (GFP) expression with the auxin-inducible DR5 promoter, containing this motif, in *Chlamydomonas reinhardtii* to prove the lack of existence of functional ARF orthologues in microalgae. While deletion analysis has shown that the canonical AuxRE is required for auxin inducibility of ARF-regulated genes, several non-canonical AuxREs have been identified (Walcher and Nemhauser 2012). These non-canonical AuxREs were determined *in silico* by a motif enrichment analysis of the promoter region of auxin induced genes in *A. thaliana*, and as such it is not known whether their relationship to auxin activity is mediated by ARFs or alternative factors (Mironova et al. 2014). Interestingly, several of these motifs are detectable in *Chlorella* sp. BR2 orthologues of genes that are induced by ARFs in land plants, such as *PATHOGENESIS-RELATED 1* and *GH3*. Further experiments will reveal whether these elements are bona fide AuxREs and their respective interactions with transcription factors.

Another important finding of this study is the discovery of the *Chlorella* orthologue of ORANGE and the functionally identical ORANGE-LIKE. This protein and its effect on carotenoid accumulation was first observed in orange cauliflower, where it was found that a mutation in the coding sequence for *ORANGE* distinguished orange cauliflower from the wild-type (Lu et al. 2006). Subsequent studies in sweet potato established that ORANGE is a DnaJ holdase, which functions by stabilizing the PSY protein, thus increasing steady-state levels of phytoene (Park et al. 2016). Evidence in the literature suggests that *ORANGE* is capable of increasing the amount of accumulated carotenoids without upregulating steps in the carotenoid biosynthetic pathway, by promoting the differentiation of plastoglobules and creating a metabolic reservoir for carotenoids (Berman et al. 2017). These attributes make *ORANGE* an interesting target for metabolic engineering, and accordingly, it would be beneficial to clone the upstream promoter sequences to better understand the regulatory factors governing its expression. Results from the transcription factor analysis may provide leads when examining proteins that bind to promoter regions of genes of interest.

### Comparison of auxin signaling in *Chlorella* sp. to other organisms

This study identified putative components of an auxin signaling network in *Chlorella* sp. BR2, including genes linked to biosynthesis, metabolism, perception and transport (Supplementary Table 4). In plants, auxin is synthesized through Trp-dependent and Trp-independent pathways. In the Trp-dependent pathway, tryptophan is synthesized from indole-3-glycerolphosphate (I3GP), and then converted into several intermediates, including tryptamine (TAM), indole-3-acetaldoxime (IAOx), indole-3-acetamide (IAM), indole-3-pyruvic acid (IPA), and then eventually IAA by the action of pathway-specific enzymes (Zhao 2012). In the Trp-independent pathway, indole, from I3GP is directly converted to IAM through *TRYPTOPHAN 2 MONOOXYGENASE* (*iaaM*), which has been shown to exist functionally in plants, but no candidate genes have been identified outside of bacterial lineages (Zhao 2012). A schematic of these pathways and the respective orthologues identified in this study are provided in Supplementary Figure 3. With this data in hand, we analysed literature reports for the presence of intermediates in the various IAA biosynthetic pathways in order to determine pathways that were likely candidates for IAA synthesis in *Chlorella* sp. BR2.

IAA has been ubiquitously detected across many clades of algae, including in 24 microalgal species (Stirk et al. 2013). The authors claimed that IAA and its precursor, IAM were the only detectable auxins in all species tested, including three species of *Chlorella*. However, the publication is lacking clarity as to whether additional auxin intermediates were quantified. A more recent study of 20 microalgal species, including 10 chlorophytes found that most species contained IAA, 2-oxindole-3-acid acid (oxIAA), IAA-Asp and IAM (Žižková et al. 2016). Further, profiling of *Chlorella minutissima* confirmed the existence of these four compounds and the precursor TAM (Stirk et al. 2014). In order to mediate auxin homeostasis, cells possess a mechanism for eliminating excess biologically active free IAA through irreversible oxidation to oxIAA or reversible conjugation with glucosinolate or amino acid groups (LeClere et al. 2002). oxIAA was found to be the key metabolite in microalgae using [^13^C]IAA, with IAA-Asp consistently falling below the limit of detection (Stirk et al. 2013; Žižková et al. 2016).

Interestingly, our transcriptome data and the genetic analysis by De Smet and Beeckman (2011) predicted the presence of the IPA pathway in *Chlorella*, which appears to be lacking based off biochemical data. It is therefore unlikely that these genes perform the same biochemical function in *Chlorella* sp. BR2. However, one study observed that IPA is rapidly non-enzymatically oxidized in solution, requiring specialized derivatization conditions to produce satisfactory measurements, which raises the possibility that previous reports *in alga* may not have detected IPA due to methodological limitations (Tam and Normanly 1998). Further experiments, involving isotopic feeding assays in conjunction with knock-downs of putative auxin biosynthesis genes, will allow for validation of putative auxin biosynthetic pathways.

This biochemical evidence, when taken in concert with our auxin-responsive transcriptome and previously published genetic data, indicates that *Chlorella* sp. BR2, and by extension, other chlorophytes, contain the machinery for at least the TAM, IAOx, IAM, indole-3acetonitrile (IAN) routes of auxin biosynthesis (De Smet and Beeckman 2011). Our data also identified putative orthologues for *GH3* family member genes, which conjugate auxin to Asp; however, no orthologues of *DIOXYGENASE FOR AUXIN OXIDATION 1*, which produces oxIAA were detected. Given the consistent detection of oxIAA as the primary IAA metabolite, it is likely that another enzyme with dioxygenase activity performs the same functions. It remains to be known whether the Trp-independent pathway of IAA synthesis exists in *Chlorella* sp. BR2, as only orthologues of *AMIDASE 1*, which catalyzes the conversion of IAM to IAA, were detected. IAM is produced by the Trp-dependent and Trp-independent pathways and as such cannot be used to deduce the existence of Trp-independent IAA biosynthesis.

An important feature of canonical auxin signaling is polar auxin transport, where the asymmetrical distribution of auxin transporters creates gradients of the hormone across individual cells and entire organs. Functional equivalence of this was explored by feeding unicellular *Chlorella vulgaris* and multicellular *Chara vulgaris* radiolabeled IAA with and without the auxin efflux inhibitor NPA. It was found that while auxin uptake in *Chara vulgaris* was facilitated by a saturable transporter, *Chlorella vulgaris* displayed a pH-dependent uptake rate which could be competitively inhibited by unlabeled IAA, indicative of a diffusion-based transport mechanism (Dibb-Fuller and Morris 1992). This study and previous genomic work found no likely orthologues of the auxin efflux PIN proteins in the genomes of eight chlorophytes, including two *Chlorella* species (De Smet and Beeckman 2011). In contrast to the physiological evidence, this study and De Smet and Beekman’s data identified putative homologues for the *AUX1/LAX1* auxin influx transporters, although the level of homology and conservation of functional residues was minimal. *PIN* proteins and *AUX1/LAX1* transporters have been characterized in *Chara corallina*, implying that polar auxin transport may be an innovation of multicellular organisms, and that unicellular orthologues of *AUX1/LAX* may perform alternative functions (Boot et al. 2012). One report observed the excretion of IAA into the culture medium by axenic *Chlorella pyrenoidosa* and *Scenedesmus aematus* cultures, but did not probe the kinetics and pH dependence of the process (Mazur et al. 2001). A strong homologue for the *ATP BINDING CASETTE B2*/*PGLYCOPROTEIN* was detected in this study for *Chlorella* sp. BR2, and other chlorophytes by De Smet and Beeckman (2011). In plants, this protein functions as an auxin efflux transporter, with knockdown mutants displaying auxin-deficient phenotypes and hypersensitivity to exogenous auxin (Cho and Cho 2013). *ABC* transporters display a uniform distribution within the plasma membrane, in contrast to the polar organization of PINs, which is indicative of a non-directional, basal auxin export role (Cho and Cho 2013). The role of a non-polar auxin efflux transporter could be imagined in the context of the colonial nature and lack of a complex body plan in unicellular chlorophytes. Putative intercellular signaling mechanisms in these organisms would likely relay environmental or metabolic information to neighbouring cells, rather than developmental cues. This would necessitate the omnidirectional transduction of signals, as the message is equally beneficial to neighboring cells in all planes. At least one member of the *ABC* family has been shown to exhibit reversible auxin efflux and influx activity, based on intracellular IAA levels, leaving open the possibility for an *ABC* orthologue with auxin influx capabilities in *Chlorella* sp. BR2 (Kamimoto et al. 2012). This could be further investigated using the specific ABC inhibitor, 2-[4-(diethylamino)-2-hydroxybenzoyl] benzoic acid (BUM) (Kim et al. 2010).

The primary means of auxin perception in land plants is through the nuclear F-box family proteins TIR1 and AFB, which form substrate-specific components of a SCF ubiquitin ligase complex (Wang and Estelle 2014). As auxin binds to SCF^TIR1/AFB^, it induces a conformational change which stimulates binding to and ubiquitination of AUX/IAA repressor proteins. These proteins are inhibitors of *ARF* transcription factors, however upon ubiquitination this repression is released (Chapman and Estelle 2009). To date, no study has been published identifying strong orthologues for AUX/IAAs and ARFs in lineages earlier than the mosses, raising questions regarding how microalgae transduce auxin perception into the now obvious transcriptional changes (De Smet and Beeckman 2011; Paponov et al. 2009). In *Klebsormidium nitens*, which lacks orthologues of the TIR1-Aux/IAA-ARF pathway a recent study reported the auxin-inducible expression of hundreds of genes, including a TF which did not change mRNA levels in the presence of cyclohexamide (Ohtaka et al. 2017). This is suggestive of an early auxin signaling process independent of SCF^TIR1/AFB^-mediated proteolysis. The Aux/IAA and ARF protein families have many paralogues in land plants, which display tissue-specific expression dynamics (Sun et al. 2015). These auxin effectors likely arose with the emergence of plants and the associated requirement for complex body plans, and as such may not be representative of a primitive system present in earlier lineages.

De Smet and Beeckman (2011) concluded that chlorophytes do not contain orthologues of *ARFs* from their genomic evidence and the observation that *Chlamydomonas reinhardtii* expressing the auxin-responsive promoter DR5 (containing TGTCTC tandem repeats) did not display IAA-inducible transcription. Lack of expression of DR5 is not sufficient evidence to rule out the existence of AuxREs and TFs in chlorophytes altogether, as AuxREs with vastly different sequences may have arisen through evolution which perform equivalent functions. Indeed, certain genes in our promoter analysis possessed a number of non-canonical AuxRE motifs. It has also been observed that some auxin-responsive genes do not contain a canonical AuxRE, allowing for the existence of alternative AuxREs in primitive lineages. This may explain why the transcriptional machinery failed to express the DR5 constructs. To this end, it would be prudent to undertake deletion assays of promoter regions for auxin-responsive genes in *Chlorella* sp. BR2 to clarify whether AuxREs do indeed exist in this species and if their structures are similar to higher plants.

Over the years, additional bona fide auxin receptor-encoding genes have been discovered in land plants, including *AUXIN BINDING PROTEIN 1* (*ABP1*), *INDOLE-3-BUTYRIC ACID RESPONSE 5* (*IBR5*) and *S-PHASE KINASE ASSOCIATED PROTEIN 2A* (*SKP2A*) (Jurado et al. 2010; Sauer and Kleine-Vehn 2011). ABP1 in plants is localized to the endoplasmic reticulum (ER) and the extracellular matrix, where it negatively regulates the endocytosis of PIN efflux transporters, and has also been implicated in the promotion of cell expansion through interaction with COP1, expansins and the regulation of gene expression through as-yet-known mechanisms (Scherer 2011). IBR5 is a nuclear-localized dual-specificity phosphatase which has been shown to physically interact with MITOGENACTIVATED PROTEIN KINASE 12 (MAPK12), where it is dephosphorylated by IBR5 in the presence of auxin, releasing auxin-responsive genes from repression (Powers and Strader 2016). Loss-of-function *ibr5* mutants show perturbed transcriptional and physiological responses to auxin in plants, which are independent of, but additive to most TIR1-mediated AUX/IAA degradation responses (Monroe-Augustus et al. 2003; Strader et al. 2008). Interestingly, a reduction in DR5 reporter expression was also observed in *ibr5* lines upon auxin treatment, suggesting cross regulation with ARFs. This provides support for the possibility that *IBR5* and *MAPK12* function in the nucleus in *Chlorella* sp. BR2 to enact auxin induced transcriptional changes in a manner that bypasses the requirement for a functional orthologue of *TIR*/*AFB*.

It would be insightful to analyse the transcriptional impact of *ibr5* and *mapk12* null mutants in microalgae to determine whether auxin responses are perturbed. MAPK signaling has also been implicated in several responses in plants, which are also seen upon auxin application in microalgae, including abiotic stress tolerance and increases in ROS detoxification activity (Jammes et al. 2011; Nowak et al. 1988; Sinha et al. 2011). Further, MAPK signaling was shown to be important in desiccation and high light stress responses in the tidal chlorophyte species *Ulva rigida* and *Chaetomorpha aerea* (Parages et al. 2014). It will be useful to study whether these responses can be perturbed by modulating *IBR5*.

SKP2A is a nuclear localized F-box protein shown to bind auxin and promote the proteolysis of E2FC and DPB cell cycle inhibitors (del Pozo et al. 2007). An *Arabidopsis SKP2A* knockdown mutant displayed an over accumulation of E2FC and DPB proteins while overexpression of the gene decreased E2FC and DPB levels, which promoted tolerance to osmotic stress challenge in *Arabidopsis* through promotion of cell division (del Pozo et al. 2007; Jurado et al. 2008). Information is lacking as to whether E2FC and DPB are the only targets of the SCF^SKP2A^ complex and the scientific community awaits a comparative transcriptome analysis with a null mutant.

### Reconstruction of putative auxin signaling pathway based on transcriptomic evidence

De Smet and Beeckman (2011) identified putative orthologues of *ABP1, IBR5* and *SKP2A* in all chlorophytes in their study, and proposed that these pathways were candidates for alternatives to the canonical SCF^TIR1^-Aux/IAA signaling in microalgae. This study extended these findings by examining the expression of these genes and other orthologues of auxin-responsive genes in land plants (Figure 9). We found that *ABP1* expression was upregulated with auxin treatment, echoing a finding by Hou et al. (2006) in *Eucommia ulmoides*. Furthermore, an alignment of our putative *ABP1* transcript with *AtABP1* revealed a moderate level of homology (40%), but a conservation of residues at sites determined to be critical for auxin binding through mutagenesis (Supplementary Figure 4). Interestingly, our predicted amino acid sequence for ABP1 only showed 45% identity to the equivalent sequence from the *Chlorella vulgaris* genome (NCBI: XP_005850997.1), with conservation of the KQEL C-terminal motif and the auxin binding residues.

**Figure 9.**
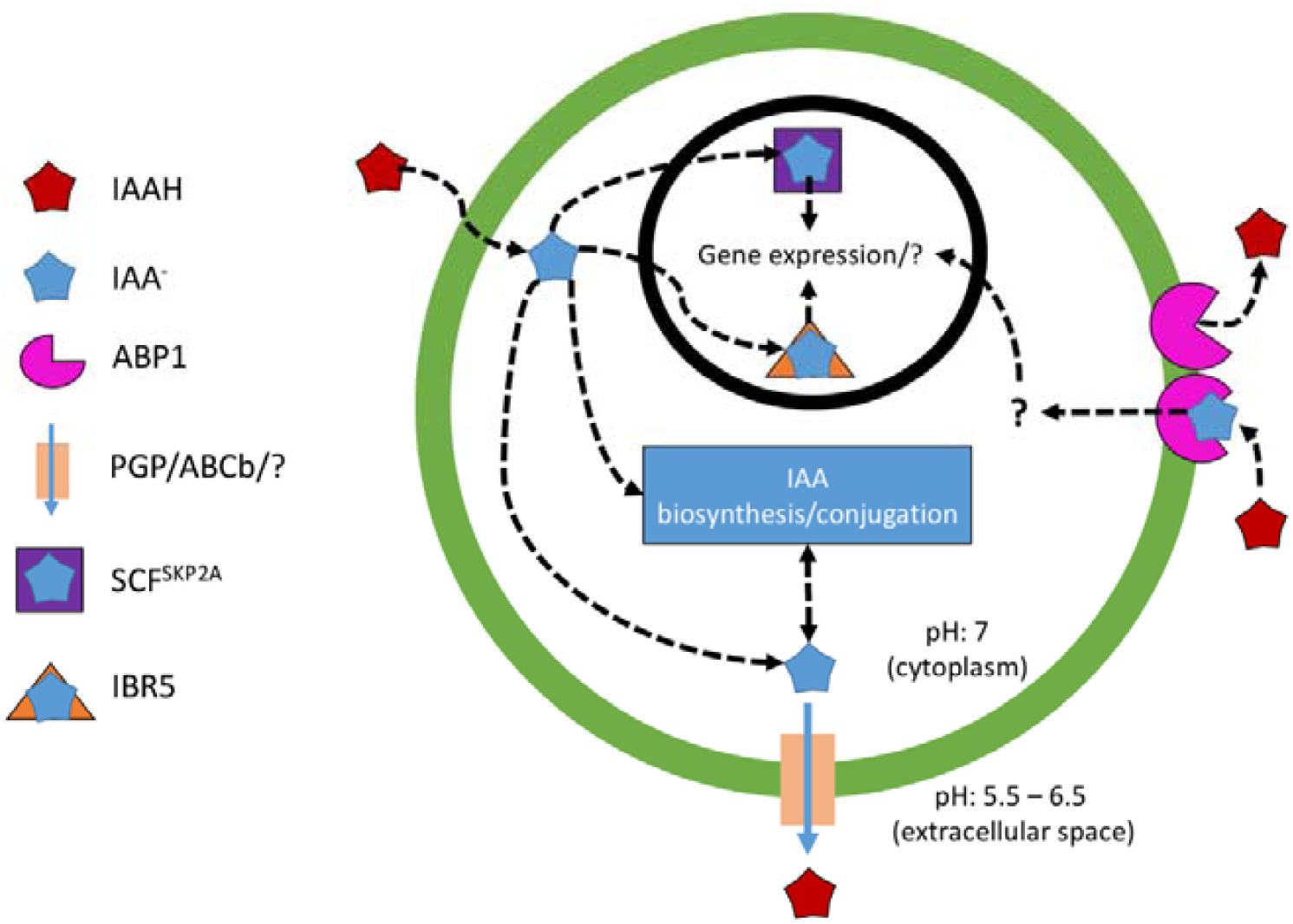
Putative model for auxin signaling in *Chlorella* sp. BR2, reconstructed from transcriptomic evidence in this study.

Interestingly, the majority of algal *ABP1* orthologues lack the C-terminal KDEL ER retention signal, present universally in plants. The relationship of this retention signal to ABP1 function remains to be elucidated, and interestingly, several mosses, including *Ceratodon purpureus, Funaria hygrometrica* and *Physcomitrella patens* only contain *ABP1* orthologues lacking the KDEL signal (Panigrahi et al. 2009). The opinion of the literature is that ABP1 levels are in constant flux through endosomal trafficking between the ER and the plasma membrane, involving a mechanism which masks the KDEL signal. ABP1 is unable to bind auxin at pHs found in the ER, while binding occurs at the relatively acidic extracellular pH. This exchange system may be a way of fine-tuning ABP1-mediated responses in more sophisticated organisms and may not be required in earlier lineages such as unicellular algae.

## Conclusion

Phytohormone-induced carotenoid production in *Chlorella* sp. BR2 was only observed in response to auxin treatment. To our knowledge, this is the first study to produce an auxin-responsive microalgal transcriptome which provides further insight into the inner workings of auxin signaling networks in unicellular microalgae. Putative metabolic targets (e.g. ABP1, SCF^SKP2A^, IBR5, ORANGE, carotenoid biosynthesis components) which may increase productivity of higher value microalgal products were also identified with the use of the assembled transcriptome.

## Acknowledgements

We wish to thank Prof. Priti Krishna, and Drs. Kent Fanning, Cristiana Dal’Molin and Faruq Ahmed for useful discussions. We are also grateful to Meat and Livestock Australia (B.NBP.0695), Australia’s Cooperative Research Centre-Project scheme (CRC-P50538) and the Saudi Arabian Government PhD scholarship program for financial support.

## Supplementary Material

**Supplementary Table 1.** Primer sequences used in qRT-PCR.

**Supplementary Table 2.** Detection and differential expression of transcripts related to carotenoid metabolic processes. DE represents the number of differentially expressed transcripts belonging to a given annotation.

**Supplementary Table 3.** Gene expression of transcripts related to antioxidant enzymes in *Chlorella* sp. BR2. NA indicates that DESeq2 was unable to infer differential expression due to undetectable expression in at least 1 replicate.

**Supplementary Table 4.** Putative orthologues of auxin biosynthesis, conjugation, transport and perception systems. Transcripts are considered as differentially expressed if their FDR is below 0.05. NA indicates that DESeq2 was unable to infer differential expression due to undetectable expression in at least 1 replicate.

**Supplementary Figure 1.** Distribution of enriched Plant GO Slim terms in upregulated transcripts of *Chlorella* sp. BR2 (FDR < 0.05).

**Supplementary Figure 2.** Distribution of enriched Plant GO Slim terms in downregulated transcripts of *Chlorella* sp. BR2 (FDR < 0.05).

**Supplementary Figure 3.** Putative auxin biosynthetic pathway for *Chlorella* sp. BR2 reconstructed from RNA-Seq data and literature reports. Grey boxes with grey text indicate enzymes lacking detectable orthologues, while grey boxes with black text show enzymes with orthologues detected in our study. Red boxes indicate metabolites suggested not to be present by literature reports. Green boxes indicate metabolites detected in previous studies in chlorophytes. Yellow boxes indicate metabolites with ambiguity in the literature regarding their presence in chlorophytes.

**Supplementary Figure 4.** Pairwise Geneious alignment with default settings of putative *Chlorella* sp. BR2 ABP1 protein (ChlABP1) with AtABP1. Black shaded areas indicate regions of similarity between the transcripts. Red rectangles indicate zinc ion-binding sites in AtABP1, while blue rectangles indicate sites shown to be involved with auxin binding in AtABP1.

